# Potential impact of novel diagnostics and treatments on the burden of antibiotic resistant in *Escherichia coli*

**DOI:** 10.1101/052944

**Authors:** Pierre Nouvellet, J.V. Robotham, N.R. Naylor, N. Woodford, Neil M. Ferguson

## Abstract

The rising threat of antibiotic resistance in Europe and beyond is of increasing concern and is prompting renewed effort to better understand and mitigate their impact. *Escherichia Coli* blood stream infections are a more major concern in Europe given their incidence and severe associated outcomes. Additionally the level of 3^rd^ generation cephalosporins and carbapenems resistance among those bacteraemia has significantly increased, limiting available treatment options. We estimated the current burden associated with *E. coli* blood stream infections in Europe at 17,000 (95%CI [8,000; 30,000]) excess deaths and 960,000 (95%CI [600,000; 1,450,000]) extra hospital bed days. From those, the contribution due to 3^rd^ generation cephalosporins and carbapenems resistant strains reached 6,000 (95%CI [2,000; 12,000]) excess deaths, and 200,000 (95%CI [76,000; 420,000]) extra hospital bed stay. In the worst case scenario, we estimated the burden of *E. coli* blood stream infection in 2026 could increase over 4-fold, mostly resulting from an increase in the level of resistance rather than an increase in the incidence of blood stream infections. Finally, we estimated that the impact of combined novel diagnostics and treatments could substantially reduce the excess mortality by 18.5% to 55.5%, and length of stay by 13.2% to 75.6%.

## Introduction

In the European Union (EU) and beyond, the emergence and rapid spread of antibiotic resistance is increasingly putting pressure on already heavily burdened health systems. In such context, predicting future trends in antibiotic resistance, its impact on populations and health systems and the potential impact of novel diagnostics and treatment with novel antibiotics is critical to plan health care capacity, establish research priorities and determine optimal interventions and investments to eventually mitigate the situation.

However, considerable uncertainty still surrounds the mechanisms, drivers and dynamics of emergence, establishment and spread of antibiotic resistance in bacterial infection (Gupta et al., 2011). While mechanistic models of resistance dynamics exists (Austin and Anderson, 1999, Lipsitch et al., 2000, Spicknall et al., 2013), current uncertainties makes their application to specific systems problematic. For instance, the diversity in bacterial species, antibiotic resistance profiles, and setting all have considerable impact on the specific dynamics of resistance. Therefore while the antibiotic surveillance systems across Europe, and indeed beyond, have improved dramatically over the last decade, it is still difficult to understand and predict the observed fluctuations both spatially and temporally. Drivers such as antibiotic consumption, importations driven by migrations, human feeding habits, and antibiotic use in the animal food industry have all been shown to play a role (Department of Health, 2014, Johnson et al., 2007, Tadesse et al., 2012, Holmes et al., 2015). However, disentangling the impact of each driver and influence factor is likely to be highly context-dependent, and will require considerable research and carefully planned experimental designs.

Where such high uncertainty exists, modelling approaches that explore scenarios can be useful in gaining better intuition on the burden, its drivers and the implications of plausible future scenarios.

In the EU, blood stream infections (BSI) caused by *Escherichia Coli* (*E. coli*) are a growing concern due to their rapidly increasing incidence (de Kraker et al., 2011a). Additionally, they affect vulnerable populations disproportionately (e.g. infants and elderly) and are associated with increased burden in terms of both severe outcomes and hospital stays (e.g. length of stay, LoS) (de Kraker et al., 2011a). Treatment options are limited, and worryingly the levels of resistance to many antibiotic classes, including fluoroquinolones, third generation cephalosporins and carbapenems (the last line treatment options), have increased over the last decades amplifying their impact (eCDC, 2015, Day et al., 2016) on both patients outcomes and health systems. With few antibiotics in the development pipeline, which would be active against such infections, the potential threat to population health is exacerbated.

The effect (and potential onward spread) of resistant organisms could be mitigated through timely, appropriate treatment. This may be afforded, for example, by novel diagnostics, enabling early identification of the causative organism, or novel antimicrobial options to which the common causative agents are not resistant.

The purpose of this work, in the context of *E. coli* associated BSI in the EU, was (a) to estimate the current burden of antibiotic resistance and its potential future trends and (b) to evaluate the potential impact of novel diagnostics and treatment options.

## Methods

We used a scenario-based approach to gain insight into the current and potential future burden of *E. coli* BSI in Europe and the potential impact of novel diagnostics and treatments. Our modelling approach consisted of four steps:

- Using surveillance data, characterise the current level of *E. coli* BSI incidence and resistance within the EU, as well as plausible future trends.
- Obtain estimates from the published literature of the per-patient excess mortality and LoS due to *E. coli* BSI and the contribution of resistance third generation cephalosporins and carbapenems.
- Using the above, evaluate the current and potential future burden of resistant infections (for hospitals), at the population level and across the EU in term of excess mortality and hospital LoS.
- Re-evaluate the outcomes of each scenario, assuming availability of new diagnostics and treatments.

### 1. Levels of E. coli BSI incidence and resistance

We first obtained the trends in *E. coli* BSI in England from the 2015 ESPAUR Report (Public Health England, 2015), as these are among the most complete data in the EU. We fitted a linear regression model to obtain the predicted current and future trend and prediction interval.

The levels of resistance to both third generation cephalosporins and carbapenems in *E. coli* and *K. pneumoniae* were explored using publicly available data from the EU EARS-net network (eCDC, 2016). These data formed the basis of the various scenarios explored:

1. Scenario 1 estimated the burden in 2016 assuming the 2016-levels of resistance would reflect the country specific observations recorded from the last 10 years among *E. coli* BSI.
2. Scenario 2 estimated the burden in 2026 assuming the 2026-levels of resistance would reflect pooled observations recorded from the last 10 years among *E. coli* BSI in the EU.
3. Scenario 3 estimated the burden in 2026 assuming the 2026-levels of resistance would reflect observations recorded from the last 10 years among *E. coli* BSI in the EU countries that recorded the worst average level of resistance.
4. Scenario 4 estimated the burden in 2026 assuming the 2026-levels of resistance would reflect pooled observations recorded from the last 10 years among *K. pneumoniae* BSI in the EU.
5. Scenario 5 estimated the burden in 2026 assuming the 2026-levels of resistance would reflect observations recorded from the last 10 years among *K. pneumoniae* BSI in the EU countries that recorded the worst average level of resistance.

We obtained samples for the 2016 and 2026 *E. coli* BSI incidences by sampling from the posterior distribution of predicted incidence in 2016 and 2026 (i.e. parametric bootstrapping). Samples of resistance levels, for both third generation cephalosporins and carbapenems, were taken by randomly choosing among observations from the EARS-net database relevant to the each specific scenario (i.e. non-parametric bootstrapping).

### 2. Excess mortality and LoS due to E. coli BSI and associated burden

We estimated the excess mortality and length of stay (LoS) per patient following the method outlined by de Kraker (de Kraker et al., 2011b). First, from published case control studies (Chang et al., 2011, de Kraker et al., 2011b), we obtained baseline (*P*_0_) and odds ratios (*aOR*-adjusted when possible) for mortality among patients with susceptible (S), third generation cephalosporins resistant (G3CR) and carbapenems resistant (CpR) *E. coli* bacteraemia. Typically, patients affected by a resistant strains of *E. coli* also suffer multiple additional conditions. Therefore, the baseline mortality (*P*_0_) represents the mortality among patients without bacteraemia but with similar comorbidity. Accordingly, the odds ratios are obtained from comparing mortality outcomes for patients with and without *E. coli* bacteraemia, whom have a similar comorbidity. We obtained estimates for the excess LoS in a similar manner.

The excess rate of deaths (D) and extra bed days (B) associated with BSIs caused by either S, G3C, Cp were then calculated using the following equations (from Bender and Blettner, 2002):

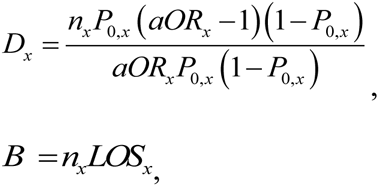

where *n*_*x*_ is the incidence of *E. coli* bacteraemia with strain *x*and *x*∊*{S,G3CR,CpR}*.

We used parametric bootstrapping to generate 100,000 samples of excess rate of deaths and extra bed days assuming a normal distribution for the log-*P* _0_‘s, log-odds’s and log-LoS’s (parameterised using the 95%CI provided in the literature).

For each scenario, using the estimate of BSI incidence and level of resistance described above, we were then able to estimate the (hospital) burden in term of both mortality and LoS. We assumed here that the population across the EU remained stable and used estimates of population sizes from the Eurostat database (Eurostat, 2016).

### 3. Potential impact of novel diagnostics and treatment

Each scenario was then re-evaluated assuming the availability of either novel diagnostics, novel treatments, or both.

We assumed that the availability of novel diagnostics would result in a reduced in time to access appropriate existing treatment(s). Therefore both the excess mortality and LoS associated with resistant-strain bacteraemias would be reduced. The outcomes for third generation cephalosporins-resistant (i.e. *aOR* and LoS) would reach a level intermediate between that currently seen (*i.e.* without novel diagnostics) and that seen for antibiotic-susceptible bacteraemias. For carbapenems-resistant bacteraemias, the reduction would be lower, reflecting the current limitations in available treatments for those infections. Thus, the outcomes (i.e. *aOR* and LoS) would reach a level intermediate between that currently seen ( *i.e.* without novel diagnostics) and that seen for third generation cephalosporins-resistant bacteraemias.

If new treatments were available, all patients could be successfully treated once accurately diagnosed. Therefore, the LoS would reach similar level as observed for antibiotic-susceptible bacteraemias after accounting for a delay in obtain appropriate diagnostics. Such delay in receiving treatment would imply that the excess mortality would be reduced to level intermediate between that currently seen and that seen for antibiotic-susceptible bacteraemias.

If novel diagnostics and treatments were both available, every cases would be immediately screened and then given a suitable treatment, tuned to the resistance level found. Therefore we assumed that the outcomes (i.e. *aOR* and LoS) would reach the level seen among patients affected by the susceptible strain

Importantly, novel diagnostics and/or treatment would not be expected to have an impact on the baseline mortality of patients.

## Results

Incidence of *E. coli* associated BSI in England (Figure 1 A) showed significant increases between 2010 and 2014 (*β*=2.8 per 100,000 per year additional cases, *p* = 0.01). As expected, for both G3C and Cp, the levels of resistance (and variability) among *K. pneumoniae* were greater than that seen among *E. coli* (Figure 1B-E). Among the worst affected countries, G3C resistance reached 40% and 100% for *E. coli* and *K. pneumoniae* respectively. For Cp resistance, those figures were lower but with similar pattern reaching 3% and over 60% for *E coli* and *K. pneumoniae* respectively.

**Figure 1:**
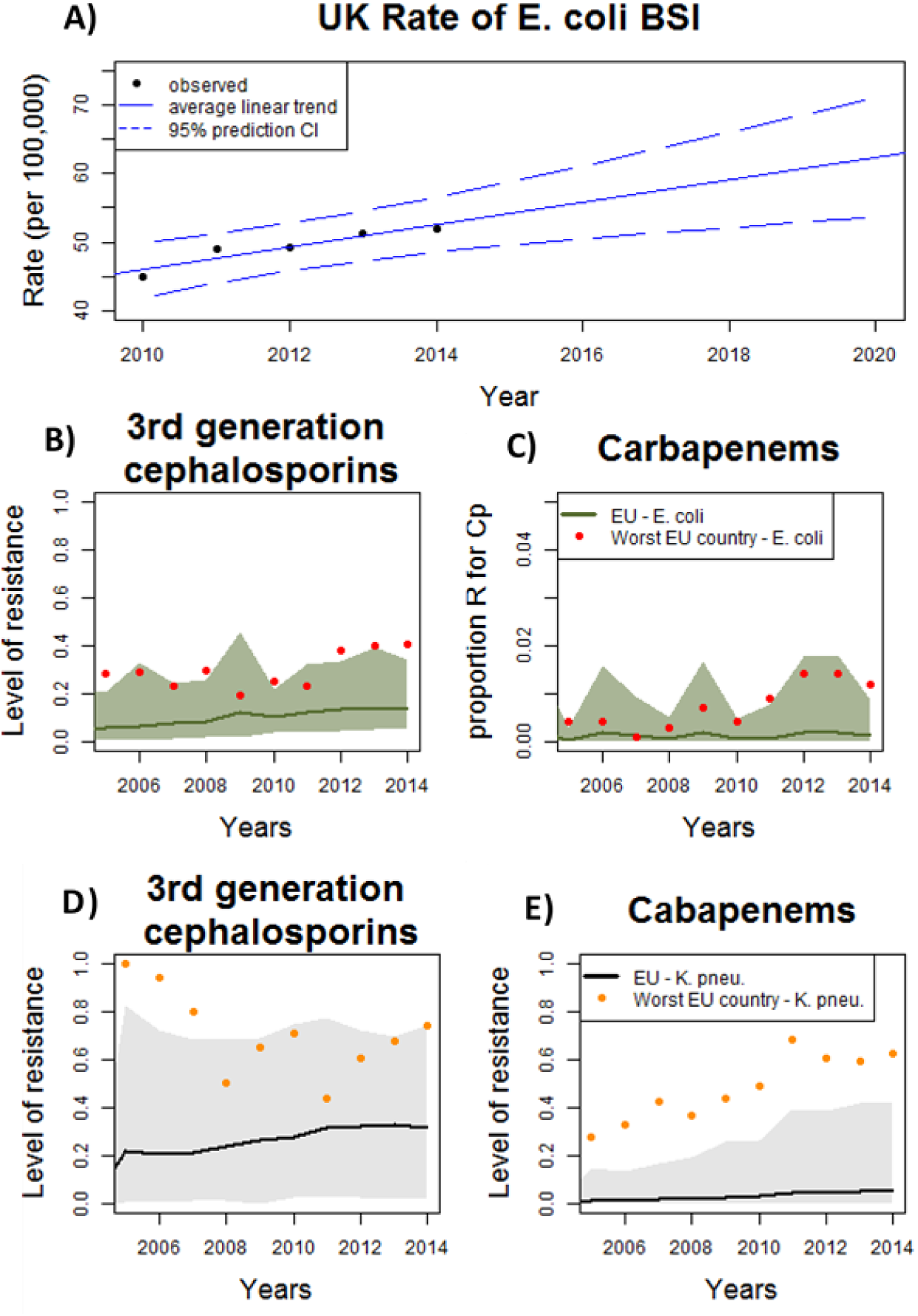
A) Trends in *E. coli*-associated BSI in the England; B) trends in the proportion of EU BSI *E-coli* isolates with third generation cephalosporins resistance; C) as B but for carbapenems resistance. The green solid line and green envelope show the mean and 95% quantile for *E. coli* in the EU. The countries sustaining the highest level of resistance are Bulgaria and Greece for *E. coli* (red dots) third generation cephalosporins and carbapenems resistance respectively. D-E) same as B-C) but showing resistance in *K. pneumoniae*. The black solid line and grey envelope show the mean and 95% quantile for *K. pneumoniae* in the EU. The countries sustaining the highest level of resistance are Romania and Greece for *K. pneumoniae* (orange dots) third generation cephalosporins and carbapenems resistance respectively.

As expected, published estimates of the baseline mortality, odd-ratio for excess mortality and excess LoS increased for patients with strain of *E. coli* that were either S, G3CR or CpR (Table 1).

**Table 1:** Baseline, excess mortality and LoS due to bacteraemia sensitive to G3C and Cp, resistant to G3C (G3CR) and resistant to Cp (CpR) among *E. coli*-associated BSI.*P*_0_ is the baseline mortality (i.e. mortality among controls with matched comorbitities but without bacteraemia) for patients with either sensitive (*S*) or resistant bacteraemia (*R*). aOR and LoS refer to the adjusted odd ratios for mortality and excess length of stay (in days) between control and case (for either sensitive or resistant bacteraemia). Here the controls are again patients without bacteraemia but with similar comorbitities. 95%CI are in parentheses. Superscripts a, b label results from De Kraker et al. (2011) and Chang et al. (2011), respectively.

| Related to sensitive bacteraemia |  |  | Related to G3CR |  |  | Related to CpR |  |  |
| --- | --- | --- | --- | --- | --- | --- | --- | --- |
| $P_{0,S}$ | $aOR_S$ | $LoS_S$ | $P_{0,R}$ | $aOR_R$ | $LoS_R$ | $P_{0,R}$ | $aOR_R$ | $LoS_R$ |
| 0.05<sup>a</sup><br>(0.01-0.09) | 1.9<sup>a</sup><br>(1.4-2.5) | 2.9<sup>a</sup><br>(1.7-4.0) | 0.55<sup>a</sup><br>(0.27-0.81) | 4.6<sup>a</sup><br>(1.7-12.3) | 7.9<sup>a</sup><br>(3.5 -<br>13.0) | 0.55<sup>a</sup><br>(0.27-0.81) | 16<sup>b</sup><br>(1.9-134.5) | 17.7<sup>b</sup><br>(3.5 -<br>65.3) |

Linking the estimated rate of BSI among the population, the level of resistance and estimates of per patient excess mortality and additional LoS attributable to infection, we obtained estimates of burden for each scenario in the absence of novel diagnostics and/or treatments (Figure 2). We estimate the current EU incidence of BSI per 100,000 of the population at 56 (95%CI: 50-61), the excess mortality at 3 deaths per 100,000 (95%CI: 1-6) and the extra bed stay due to *E.coli* bacteraemia at 186 days per 100,000 (95%CI: 116-283). Over the EU population, this implies 290,000 (95%C: 260,000-310,000) *E. coli* associated BSI occurring in 2016. Of those, we estimate 25,000 (95%CI: 16,000-34,000) to be caused by drug-resistant bacteria. We further estimate total *E. coli* bacteraemias to result in 17,000 (95%CI: 8,000-30,000) excess deaths in 2016 and in 960,000 (95%CI: 600,000-1,450,000) extra bed days. Of these, resistant strains cause 6,000 (95%CI: 2,000-12,000) deaths, and 200,000 (95%CI: 76,000-420,000) extra bed days.

**Figure 2:**
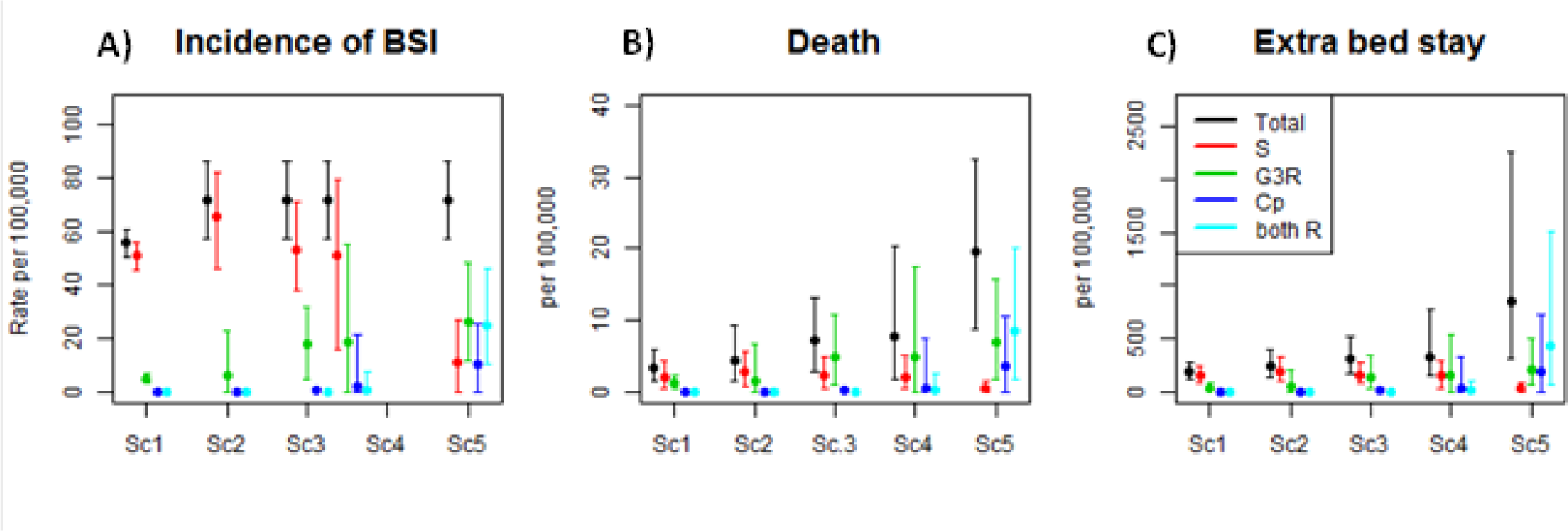
Predicted A) Incidence, B) extra deaths and C) extra day of hospital stay (per 100,000 of the general population) associated with *E. coli*-associated BSI in the EU. Graphs show overall estimates and contributions from bacteraemia sensitive to both antibiotics, resistant to either G3C or Cp, or resistant to both antibiotics. The scenarios (Sc1-Sc5) are detailed in the text.

An increase in BSI incidence alone would significantly (and linearly) increase the burden of *E. coli* bacteraemia. Comparing Sc1 and Sc2, we see a 1.3-fold increase in both BSI incidence, excess deaths and extra bed days. However, an increase in the proportion of resistant strains (Sc 3-5) could potentially have a much larger impact on the burden of *E. coli* bacteraemia; Sc5, the worst-case we examined, shows a more than 4-fold increase in both excess death and extra bed days compared with Sc2.

Finally, we predicted the impact of novel diagnostics and treatments on the hospital burden of *E. coli* BSI (Table 3). We estimate that the impact of new diagnostics alone would reduce mortality by a percentage ranging from 7.7% to 16.2%, whilst the impact of new treatments alone was estimated to range from a 10.2% to a 38.5% reduction, with greater impact when the level of resistance was higher. Comparatively, the impact on length of stay ranged from a 6.5% to a 27.6% reduction for new diagnostics (from 5.5% to 54.2% for new treatments) again with greater impact in scenarios with higher resistance levels. Generally, novel treatments were associated with a greater impact than novel diagnostics, especially for the length of stay.

**Table 3:**
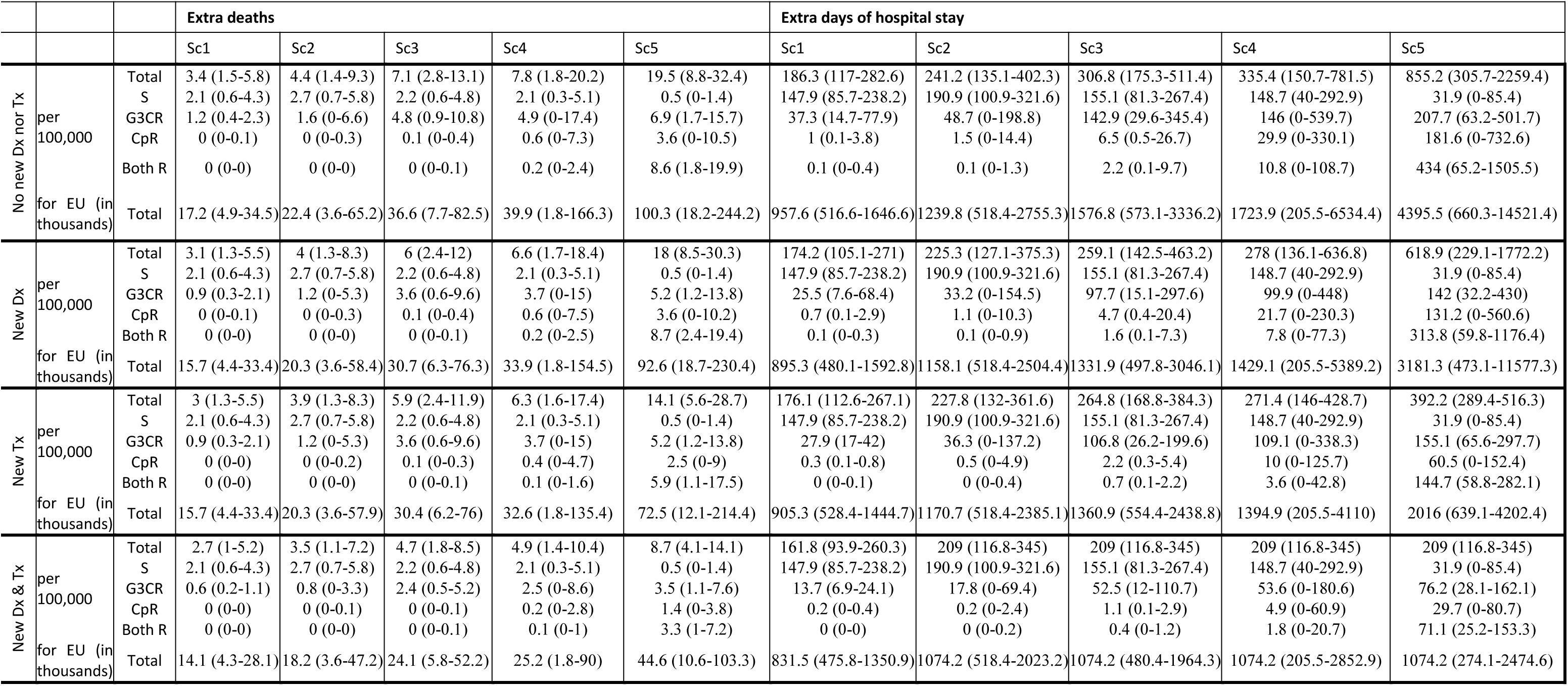
A) Incidence, B) extra deaths and C) extra days of hospital stay associated with *E. coli* BSI in the EU under each of the five scenarios (Sc1-Sc5) detailed in the text. Dx is an abbreviation for novel diagnostics and Tx for novel treatments.

The combined impact of both novel diagnostics and treatments was much greater with excess mortality reduced by 18.5% to 55.5%, and length of stay reduced by 13.2% to 75.6%. For instance, in the worst scenario (Sc. 5), the implementation of new diagnostics and treatments would reduce the excess death from 19.5 (95%CI: 8.8-32.4) to 8.7 (95%CI: 4.1-14.1) deaths per 100,000, and the excess length of stay from 855 (95%CI: 306-2260) to 209 (95%CI: 117-345) days per 100,000.

## Discussion

With close to 300,000 estimated *E. coli* associated BSI diagnosed in the EU annually, we estimated the annual burden of BSI at around 20,000 extra deaths and a million extra bed days, of which roughly a fifth could be linked to infections caused by strains resistant to third generation cephalosproins or carbapenem. These estimates are in good agreement with previous work (de Kraker et al., 2011a) accounting for increases in BSI incidence and our inclusion of the burden associated with both third generation cephalosporins and cabapenems. Worryingly, our worst-case, albeit unlikely, scenario suggests these numbers might increase 4fold in coming years.

We predicted that the impact of new treatments would be marginally greater than new diagnostics alone, but that each has a limited impact individually. However, our analysis suggests the combined impact of both would be at least additive, and in the worst-case scenario synergistic – achieving substantial reductions in disease burden (>50% in the worst-case scenario). Additionally, we found that halting the rise in incidence of BSI (e.g. through enhanced infection prevention and control for example) would have an impact on the burden of *E. coli* associated BSI comparable to the development of either new treatment or diagnostics.

While potentially informative to policy makers and to guide future research activities, our results should be cautiously interpreted. Our work aims to give intuition and examine plausible scenarios and to illustrate key gaps in current knowledge. Reliable estimates of clinical outcomes for patients affected by resistant strains remain scarce, but are critical in estimating the burden of antibiotic resistance. As argued by de Kraker et al. (2011b), a case-control study should match cases with patients with similar aetiology but without bacteraemia, thereby allowing estimates to be derived which control for heterogeneities in baseline mortality. Yet in most case-control studies, patients with a resistant strain are matched to patients with a susceptible strain, potentially inflating estimates of burden associated with resistance, since patients with resistant strains typically have greater co-morbidities. Likewise, studies ignoring the timing of infection (*i.e.* which do not match on infection time) lead to bias and often overestimation of attributable length of stay (Barnett et al., 2011, Nelson et al., 2015). Therefore, interpreting results from such experimental designs in term of excess mortality or length of stay may be misleading.

While our estimates focus on excess mortality and length of stay, other outcomes would be of interest. For instance, patients seemingly recovering from resistant strain bacteraemia could be at an increased risk of re-infection and re-admission to hospital (Bart et al., 2015). Additionally, the long-term development of co-morbidities (during and after bacterial infection) is seldom explored but could also significantly affect burden estimates for antimicrobial resistance in the hospital context.

The simple and informative approach used here illustrates the invaluable insight that can be gained from accurate surveillance systems for both disease incidence and resistance levels. While countries like the UK have considerably improved their surveillance over the last decade, data for many countries, some with a likely high burden, are still limited. Given the need for a global approach in tackling antimicrobial resistance (Levy and Marshall, 2004, Wernli et al., 2011), standardisation of surveillance would greatly benefit the prioritisation of actions, as exemplified by the ERA-net network (eCDC, 2016).

Finally, our analysis highlights gaps in our understanding of the dynamics of both bacteraemia incidence and resistance. The lack of dynamical mathematical models tested and validated to predict variations in incidence and level of resistance drove us to use simpler scenario modelling rather than the more sophisticated models commonly used to understand burden while accounting for the disease dynamics (Dye, 2012, Nouvellet et al., 2015). The key limitation of the scenario modelling presented is that it lacks a mechanistic representation of the spread of infection and resistance. Thus we are unable to capture the indirect effects of improved treatment and diagnostics on onward transmission. However, before mechanistic models can be justified in this context, better understanding the drivers of both incidence and resistance levels is needed.

Our analysis provides an assessment of the current and future burden associated with *E. coli* bloodstream infection in hospital setting. A substantial proportion of this burden was estimated to result from third generation cephalosporins and carbapenems resistant bacteraemias, illustrating the pressing need for the rapid development of novel diagnostics and treatments. In further assessing the potential impact of such novel diagnostics and treatments strategies, we demonstrate the benefit of increasing capacity for research and development in this area.

## Acknowledgements

The authors acknowledge research funding from the UK Medical Research Council and the UK NIHR under the Health Protection Research Unit initiative.

